# Transcriptional profiling of macrophages derived from monocytes and iPS cells identifies a conserved response to LPS and novel alternative transcription

**DOI:** 10.1101/014209

**Authors:** Kaur Alasoo, Fernando O. Martinez, Christine Hale, Siamon Gordon, Fiona Powrie, Gordon Dougan, Subhankar Mukhopadhyay, Daniel J. Gaffney

## Abstract

Macrophages differentiated from human induced pluripotent stem cells (IPSDMs) are a potentially valuable new tool for linking genotype to phenotype in functional studies. However, at a genome-wide level these cells have remained largely uncharacterised. Here, we compared the transcriptomes of naïve and lipopolysaccharide (LPS) stimulated monocyte-derived macrophages (MDMs) and IPSDMs using RNA-Seq. The IPSDM and MDM transcriptomes were broadly similar and exhibited a highly conserved response to LPS. However, there were also significant differences in the expression of genes associated with antigen presentation and tissue remodelling. Furthermore, genes coding for multiple chemokines involved in neutrophil recruitment were more highly expressed in IPSDMs upon LPS stimulation. Additionally, analysing individual transcript expression identified hundreds of genes undergoing alternative promoter and 3′ untranslated region usage following LPS treatment representing a previously under-appreciated level of regulation in the LPS response.

## Introduction

Macrophages are key cells associated with innate immunity, pathogen containment and modulation of the immune response^1,2^. Commonly used model systems for studying macrophage biology have centered on macrophage-like leukemic cell lines, primary macrophages derived from model organisms and primary human macrophages differentiated from blood monocytes. Although these cells have provided important insights into macrophage-associated biology, there are issues that need consideration. Immortalised cell lines often have abnormal genetic structures and can exhibit functional defects compared to primary cells^3,4^, while multiple functional differences exist between macrophages from different species^5^.

Additionally, introducing targeted genetic modifications into the genomes of human monocyte derived macrophages (MDMs) still remains challenging, limiting their utility in genetic studies. In particular, lipid based transfection methods show low efficiency in macrophages. Transduction of macrophages using virus based vectors has relatively higher efficiency but they can integrate into the host genome or impact macrophage function in other ways^6^. Finally, introduction of foreign nucleic acid into the cytosol induces a robust antiviral response that may make it difficult to interpret experimental data^7^. Undifferentiated iPS cells are relatively easier to transfect and do not suffer from the same limitations.

Recently, methods have been developed to differentiate macrophage-like cells from human induced pluripotent stem (iPS) cells that have the potential to complement current approaches and overcome some of their limitations^8,9^. This approach is scalable and large numbers of highly pure iPS-derived macrophages (IPSDMs) can be routinely obtained from any human donor following initial iPS derivation. IPSDMs also share striking phenotypic and functional similarities with primary human macrophages^8,9^. Since human iPS cells are amenable to genetic manipulation, this approach can provide large numbers of genetically modified human macrophages^9^. Previous studies have successfully used IPSDMs to model rare monogenic defects that severely impact macrophage function^10^. However, it remains unclear how closely IPSDMs resemble primary human monocyte-derived macrophages (MDMs) at the transcriptome level and to what extent they can be used as an alternative model for functional assays.

Here, we provide an in-depth comparison of the global transcriptional profiles of naïve and lipopolysaccharide (LPS) stimulated IPSDMs with MDMs using RNA-Seq. We found that their transcriptional profiles were broadly similar in both naïve and LPS-stimulated conditions. However, certain chemokine genes as well as genes involved in antigen presentation and tissue remodelling were differentially regulated between MDMs and IPSDMs. Additionally, we identified novel changes in alternative transcript usage following LPS stimulation suggesting that alternative transcription may represent an important component of the macrophage LPS response.

## Results

### Gene expression variation between iPS cells, IPSDMs and MDMs

RNA-Seq was used to profile the transcriptomes of MDMs derived from five and IPSDMs derived from four different individuals (Methods). Identical preparation, sequencing and analytical methodologies were used for all samples. Initially, we used Principal Component Analysis (PCA) to generate a genome-wide overview of the similarities and differences between naïve and LPS-stimulated IPSDMs and MDMs as well as undifferentiated iPS cells. The first principal component (PC1) explained 50% of the variance and clearly separated iPS cells from all macrophage samples (Fig. 1, Supplementary Fig. S1) illustrating that IPSDMs are transcriptionally much more similar to MDMs than to undifferentiated iPS cells. This was further confirmed by hierarchical clustering (Supplementary Fig. S2) as well as high expression of macrophage specific markers and low expression of pluripotency factors in IPSDMs (Supplementary Fig. S3). The second PC separated naïve cells from LPS-stimulated cells and explained 16% of the variance, while the third PC, explaining 8% of the variance, separated IPSDMs from MDMs. The principal component that separated IPSDMs from MDMs (PC3) was different from that separating macrophages from iPS cells (PC1). Since principal components are orthogonal to one another, this suggests that the differences between MDMs and IPSDMs are beyond the simple explanation of incomplete gene activation or silencing compared to iPS cells.

**Figure 1.**
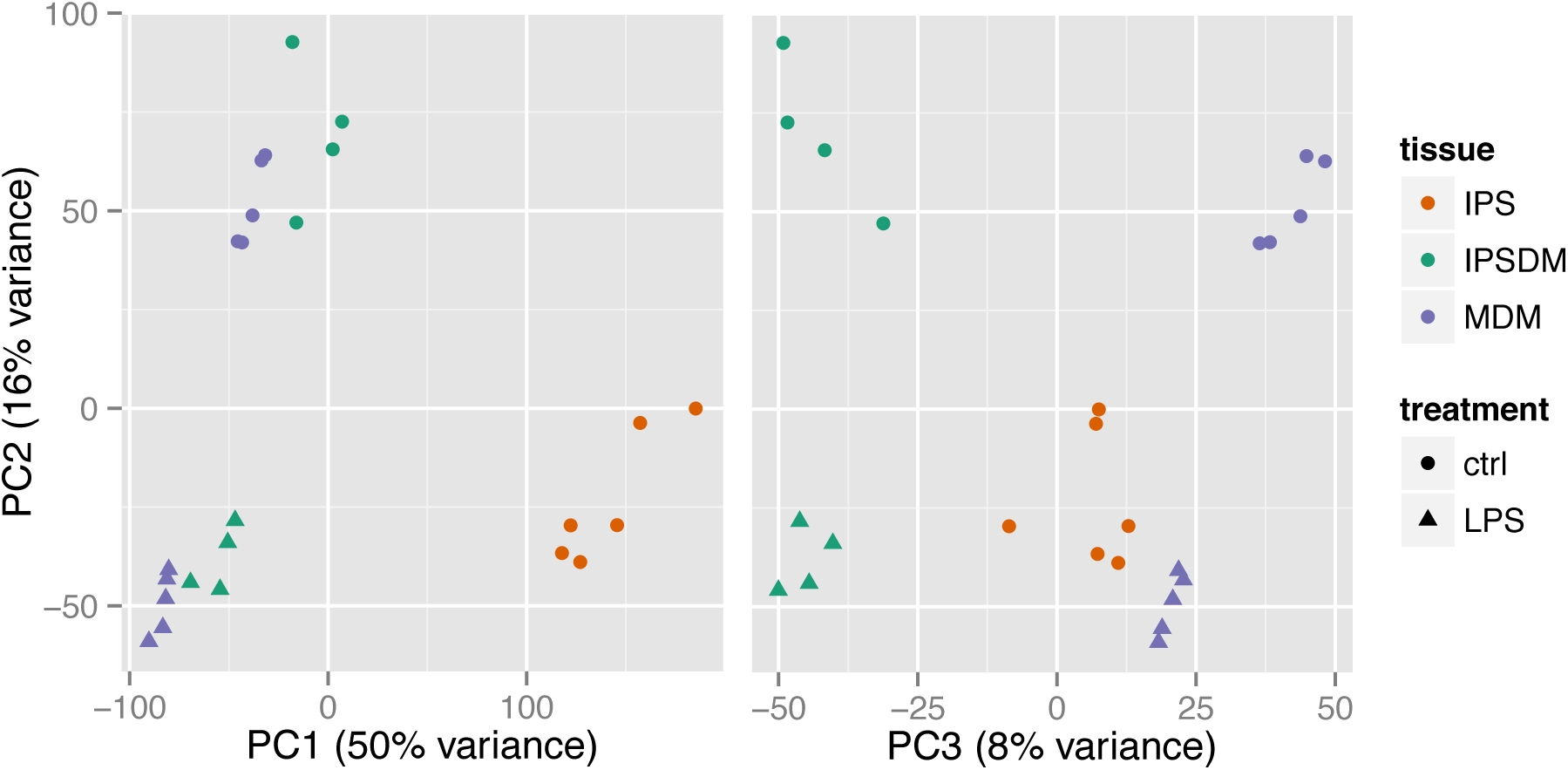
Gene expression variation between iPS cells, IPSDMs and MDMs. Principal Component Analysis of expressed genes (TPM > 2) in iPS cells, IPSDMs and MDMs.

### Differential expression analysis of IPSDMs vs MDMs

Although PCA provides a clear picture of global patterns and sources of transcriptional variation across all genes in the genome, important signals at individual genes might be missed. To better understand transcriptional changes at the gene level we used a two factor linear model implemented in the DESeq2 pacakge^11^. The model included an LPS effect, capturing differences between unstimulated and stimulated macrophages and a macrophage type effect capturing differences between MDMs and IPSDMs. Our model also included an interaction term that identified genes whose response to LPS differed between MDMs and IPSDMs. We defined significantly differentially expressed genes as having a fold-change of >2 between two conditions using a p-value threshold set to control our false discovery rate (FDR) to 0.01.

Using these thresholds, we identified 2977 genes that were differentially expressed between unstimulated IPSDMs and MDMs (Supplementary Table S1). Among these genes, 2080 were more highly expressed in IPSDMs and 897 were more highly expressed in MDMs (Fig. 2a). Genes that were more highly expressed in MDMs such as HLA-B, LYZ, MARCO and HLA-DRB1 (Fig. 2c), were significantly enriched for antigen binding, phagosome and lysosome pathways (Table 1, Supplementary Table S2). This result is consistent with a previous report that MDMs have higher cell surface expression of MHC-II compared to IPSDMs^8,9^. Genes that were more highly expressed in IPSDMs, such as MMP2, VEGFC and TGFB2 (Fig. 2c) were significantly enriched for cell adhesion, extracellular matrix, angiogenesis, and multiple developmental processes (Table 1, Supplementary Table S2).

**Figure 2.**
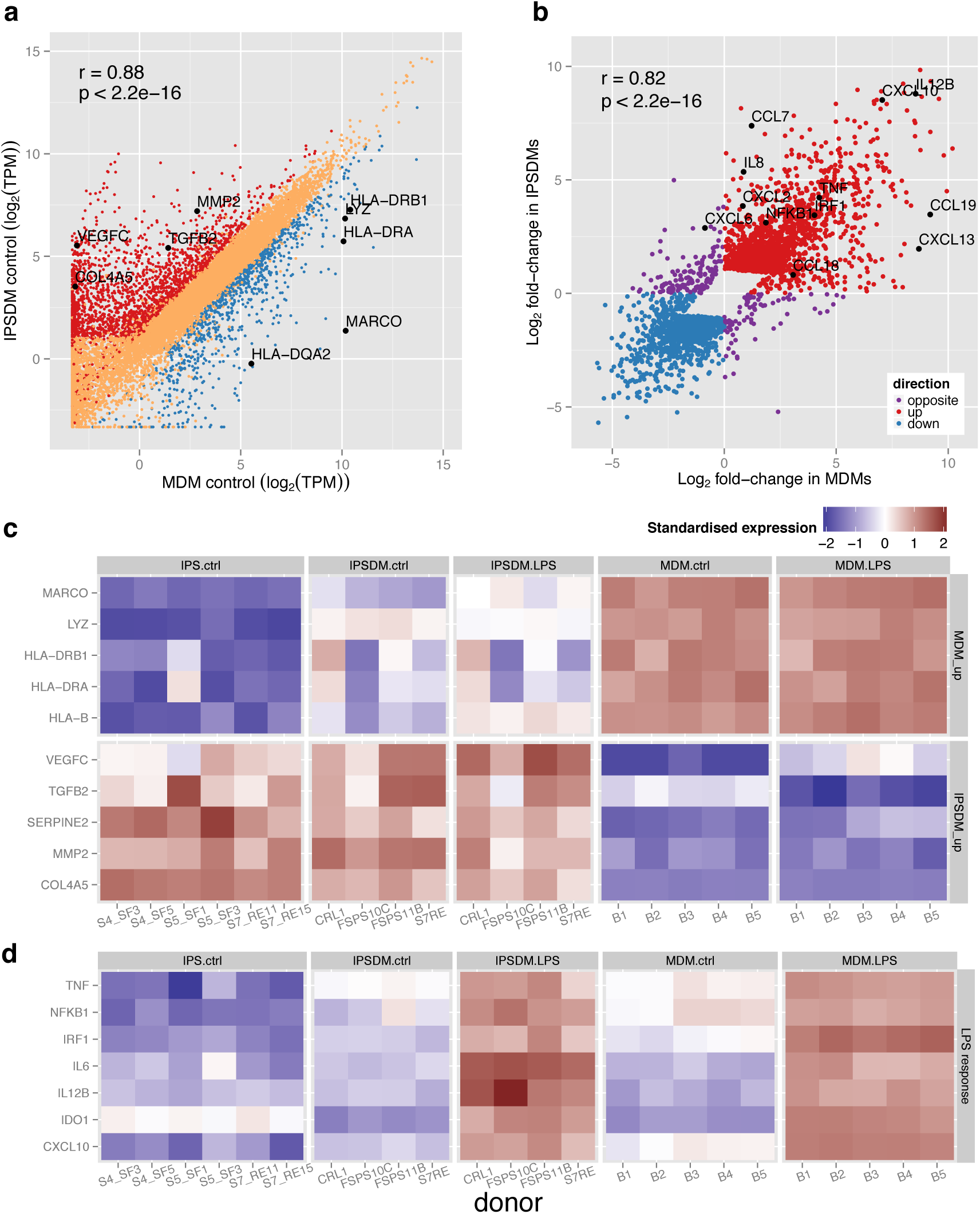
Differential expression analysis of IPSDMs vs MDMs. **(a)** Scatter plot of gene expression levels between MDMs and IPSDMs. Genes that are significantly more highly expressed in IPSDMs are shown in red and genes that are significantly more highly expressed in MDMs are shown in blue. **(b)** Scatter plot of fold change in response to LPS between MDMs (x-axis) and IPSDMs (y-axis). Only genes with significant LPS or interaction term in the linear model are shown. Genes where LPS response is in opposite directions between MDMs and IPSDMs are highlighted in purple. Raw data is presented in Supplementary Table S3. **(c)** Heat map of genes differentially expressed between MDMs and IPSDMs. Representative genes from significantly overrepresented Gene Ontology groups (Table 1) include antigen presentation (HLA genes), lysosome formation (LYZ), angiogenesis (VEGFC, TGFB2), and extracellular matrix (SERPINE2, MMP2 COL4A5). The same genes are also marked in panel a. **(d)** Heatmap of example genes upregulated in LPS response.

**Table 1.** Selection of enriched Gene Ontology terms and KEGG pathways for different groups of differentially expressed genes. Full results are given in Supplementary Table S2.

| **Upregulated in LPS response** |  |  |  |
| --- | --- | --- | --- |
| **Term ID** | **Domain** | **Term name** | **p-value** |
| GO:0045087 | BP | innate immune response | 7.31E-45 |
| GO:0009617 | BP | response to bacterium | 2.42E-28 |
| GO:0032496 | BP | response to lipopolysaccharide | 4.38E-28 |
| KEGG:04668 | ke | TNF signaling pathway | 1.71E-20 |
| KEGG:04064 | ke | NF-kappa B signaling pathway | 3.56E-14 |
| **Downregulated in LPS response** |  |  |  |
| **Term ID** | **Domain** | **Term name** | **p-value** |
| GO:0005096 | MF | GTPase activator activity | 1.01E-09 |
| GO:0007264 | BP | small GTPase mediated signal transduction | 3.14E-09 |
| **More highly expressed in MDMs compared to IPSDMs** |  |  |  |
| **Term ID** | **Domain** | **Term name** | **p-value** |
| GO:0050778 | BP | positive regulation of immune response | 1.97E-21 |
| GO:0003823 | MF | antigen binding | 2.55E-18 |
| GO:0005764 | CC | lysosome | 1.42E-17 |
| GO:0034341 | BP | response to interferon-gamma | 2.17E-16 |
| GO:0042611 | CC | MHC protein complex | 3.67E-16 |
| KEGG:04612 | ke | Antigen processing and presentation | 3.47E-13 |
| KEGG:04145 | ke | Phagosome | 2.46E-11 |
| **More highly expressed in IPSDMs compared to MDMs** |  |  |  |
| **Term ID** | **Domain** | **Term name** | **p-value** |
| GO:0030198 | BP | extracellular matrix organization | 3.05E-45 |
| GO:0016477 | BP | cell migration | 1.50E-40 |
| GO:0001568 | BP | blood vessel development | 4.89E-36 |
| GO:0016337 | BP | cell-cell adhesion | 6.27E-25 |
| GO:0001525 | BP | angiogenesis | 1.34E-24 |

In the LPS response we identified 2638 genes that were differentially expressed in both MDMs and IPSDMs, of which 1525 genes were upregulated while 1113 were downregulated (Supplementary Table S1). As might be expected, Gene Ontology and KEGG pathway analysis revealed large enrichment for terms associated with innate immune and LPS response, NF-κB and TNF signalling (Table 1, Supplementary Table S2). We also identified 569 genes whose response to LPS was significantly different between IPSDMs and MDMs (Supplementary Table S1 and S3). The majority of these genes (365 genes) were up- or downregulated in both macrophage types but the magnitude of change was significantly different and only 229 genes were regulated in the opposite direction (8.7% of the LPS-responsive genes). This set of 229 were much weaker responders to LPS overall (2.3-fold compared to 4.7-fold). Additionally, we could find no convincing pathway or Gene Ontology enrichment signals in either gene set (229 and 569 genes) compared to all LPS-responsive genes. Overall, we found that the fold change of the genes that responded to LPS was highly correlated between MDMs and IPSDMs (r = 0.82, Fig. 2b) indicating that the LPS response in these two macrophage types is broadly conserved. Interestingly, we also found that mean fold change was marginally (10%) higher in MDMs (4.95) compared to IPSDMs (4.43). The behaviour of some canonical LPS response genes is illustrated in Fig. 2d.

Although we did not identify pathways or Gene Ontology terms that were particularly enriched in differentially regulated genes, we noticed that IL8 and CCL7 mRNAs were particularly upregulated in IPSDMs compared to MDMs (Fig. 2b). Consequently, we looked at the response of all canonical chemokines in an unbiased manner^12^. Interestingly, we observed relatively higher induction of further CXC subfamily monocyte and neutrophil attracting chemokines in IPSDMs (Fig. 3). Moreover, five out of seven CXCR2 ligands^12^ were more strongly induced in IPSDMs (FDR < 0.1, fold-change difference between MDMs and IPSDMs > 2) which is significantly more than is expected by chance (Fisher’s exact test p=4.531e-06) (Fig. 3). These genes were also expressed at substantial levels (TPM > 100, Supplementary Table S4), with IL8 being one of the most highly expressed gene in IPSDMs after LPS stimulation. On the other hand, MDMs displayed relatively higher induction of three chemokines involved in attracting B-cells, T-cells and dendritic cells (CCL18, CCL19, CXCL13) (Fig. 3)

**Figure 3.**
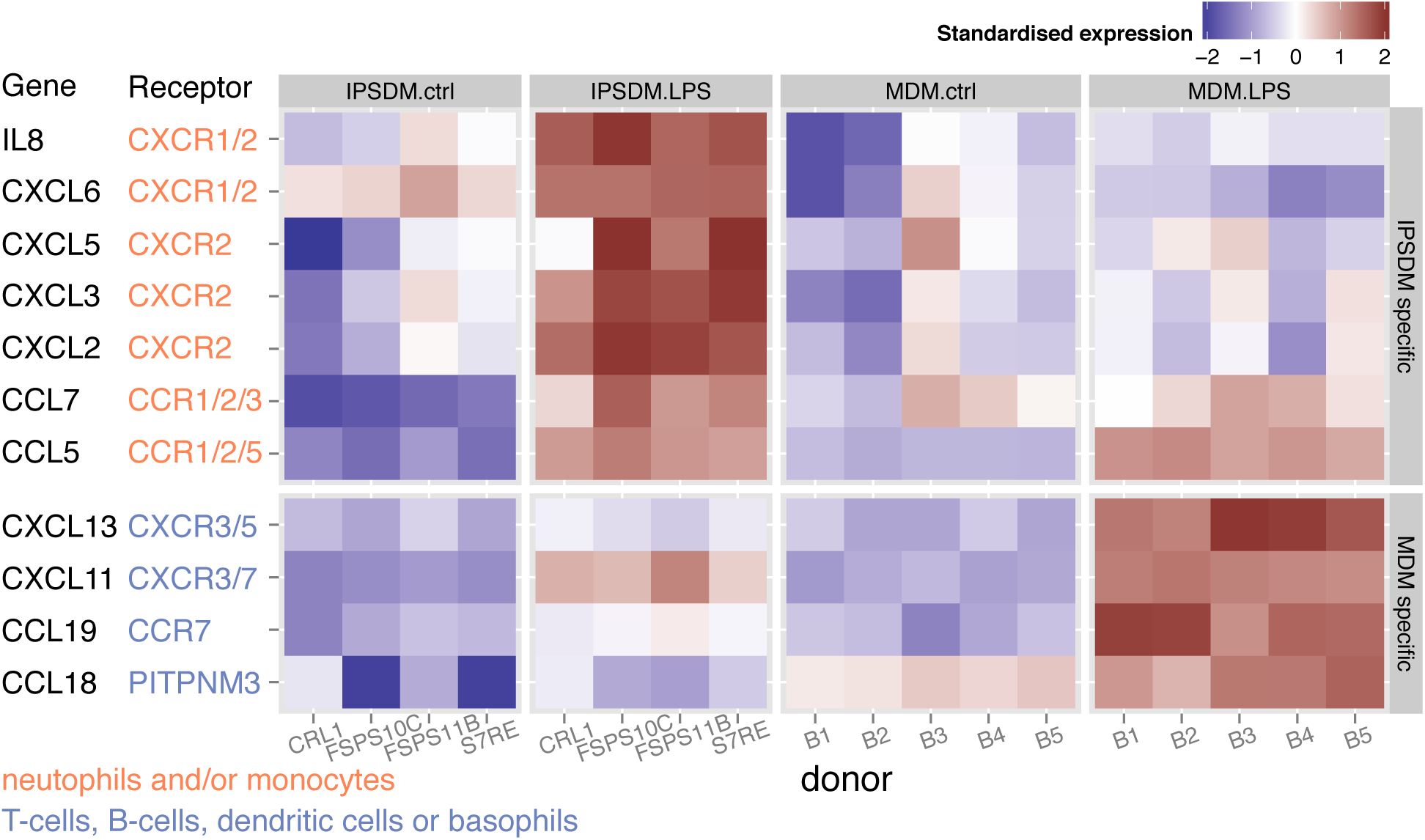
Chemokine genes that were particularly upregulated in either IPSDMs or MDMs in LPS response. Their annotated receptors and target cell types were taken from the literature^12,30^. Mean absolute expression values are shown in Supplementary Table S4.

### Global variation in alternative transcript usage

Many human genes express multiple transcripts that can differ from each other in terms of function, stability or sub-cellular localisation of the protein product^13,14^. Considering expression only at a whole gene level can hide some of these important differences. Therefore, we sought to quantify how similar were naïve and stimulated IPSDMs and MDMs at the individual transcript expression level. Here, we first used mmseq^15^ to estimate the most likely expression of each annotated transcript that would best fit the observed pattern of RNA-Seq reads across the gene. Next, we calculated the proportion of each transcript by dividing transcript expression by the overall expression level of the gene, only including genes that were expressed over two transcripts per million (TPM)^16^ in all experimental conditions (8284 genes). Since the proportions of all transcripts of a gene sum up to one and most genes express one dominant transcript^17^, we used the proportion of the most highly expressed transcript to represent the transcriptional make-up of the gene. In this context and similarly to gene level analysis, the first PC explained 31% of the variance and clearly separated IPS cells from macrophages (Fig. 4a). However, here the second PC (11% of variance) not only separated unstimulated cells from stimulated cells but also IPSDMs from MDMs. One interpretation of this result is that the changes in transcript usage between IPSDMs and MDMs, to some extent, also resemble those induced in the LPS response. Further analysis (below) highlighted that much of this variation can be explained by changes in 3′ untranslated region (UTR) usage.

**Figure 4.**
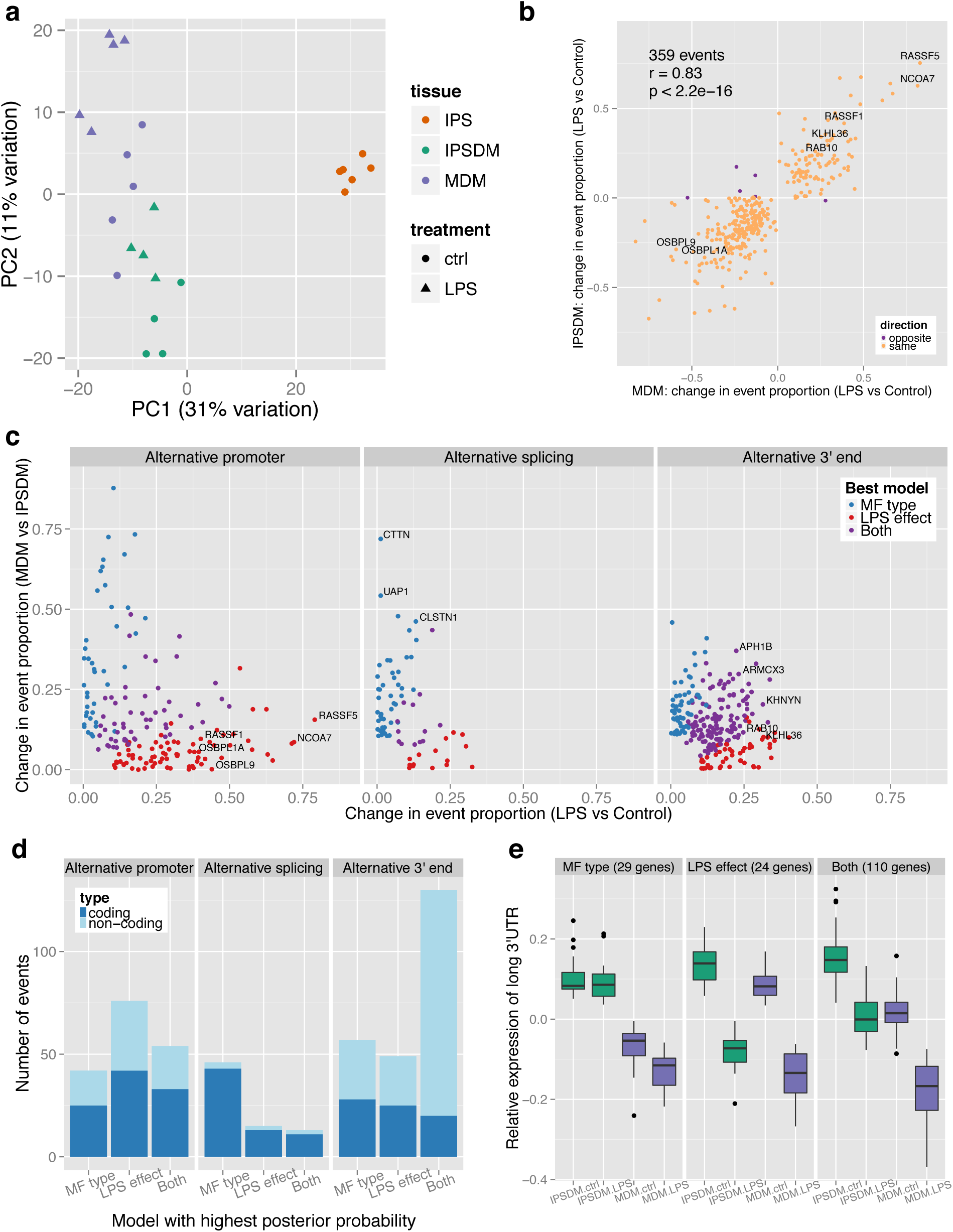
Alternative transcription in IPSDMs and MDMs. **(a)** Principal component analysis of relative transcript proportions in IPS cells, IPSDMs and MDMs. Only genes with mean TPM > 2 in all conditions were included. **(b)** Alternative transcription events detected in LPS response. Each point corresponds to an alternative transcription event and shows the absolute change in the proportion of the most highly expressed transcript (across all samples) in LPS response in MDMs (x-axis) and IPSDMs (y-axis). **(c)** All detected alternative transcription events were divided into three groups based on whether they affected alternative promoter, alternative splicing or alternative 3′ end of the transcript. For each event, we plotted its change in proportion in LPS response (x-axis) against its change between macrophage types (y-axis). The events are coloured by the most parsimonious model of change selected by mmseq: LPS effect (difference between naïve and LPS-stimulated cells only); macrophage (MF) type (difference between IPSDMs and MDMs only); both (data support both MF type and LPS effects). **(d)** Number of alternative transcription events form panel C grouped by position in the gene (alternative promoter, alternative splicing, alternative 3′ end) and most parsimonious model selected by mmseq. **(e)** Relative expression of long alternative 3′ UTRs in genes showing a change between IPSDM and MDMs (MF type), between naïve and LPS-stimulated cells (LPS effect) and for genes showing both types of change.

### Identification and characterisation of alternative transcription events

Alternative transcription can manifest in many forms, including alternative promoter usage, alternative splicing and alternative 3′ end choice, each likely to be regulated by independent biological pathways. Thus, we sought to characterise and quantify how these different classes of alternative transcription events were regulated in the LPS response, and between MDMs and IPSDMs. Using a linear model implemented in the mmdiff^18^ package followed by a series of downstream filtering steps (Methods) we identified 504 alternative transcription events (ATEs) in 485 genes (Supplementary Table S5). Out of those, 145 events changed between unstimulated IPSDMs and MDMs (macrophage (MF) type effect) while 156 events changed between naive and LPS stimulated cells across macrophage types (LPS effect). Further 197 events had different baseline expression between macrophage types, but also changed in the same direction after LPS stimulation (Both effects). Finally, only 6 events change in the opposite direction after LPS stimulation between MDMs and IPSDMs (Fig. 4b). Next, we focussed on the 359 events that changed in the LPS response in at least one macrophage type (156 + 197 events with LPS response in the same direction and 6 events with LPS response in the opposite direction). We found that the LPS-induced change in the proportion of the most highly expressed transcript was highly correlated between MDMs and IPSDMs (Pearson r = 0.83) (Fig. 4b), further confirming that the LPS response in both macrophage types is conserved. Interestingly, we found a highly significant overlap between differentially expressed and alternatively transcribed genes in LPS response (63/150, hypergeometric test p<2.4×10^-10^), but no significant overlap between differentially expressed and alternatively transcribed genes between MDMs and IPSDMs (28/140, hypergeometric test p < 0.74).

Perhaps surprisingly, although the transcriptional response to LPS at the whole gene level is relatively well understood, the effect of LPS on transcript usage has remained largely unexplored. Therefore we decided to investigate the types of alternative transcription events identified in the LPS response as well as between MDMs and IPSDMs. Most protein coding changes in the LPS response were generated by alternative promoter usage (Fig. 4c-d). In total, we identified 180 alternative promoter events, 51 of which changed the coding sequence by more than 100 bp in the LPS response (Supplementary Table S5, Supplementary Fig. S4). Strikingly, alternative promoter events displayed larger change in proportion than other events and they often changed the identity of the most highly expressed transcript of the gene (Fig. 4c). Strong activation of an alternative promoter in the NCOA7 gene is illustrated in Fig. 5a. More examples can be found in Supplementary Fig. S4.

**Figure 5.**
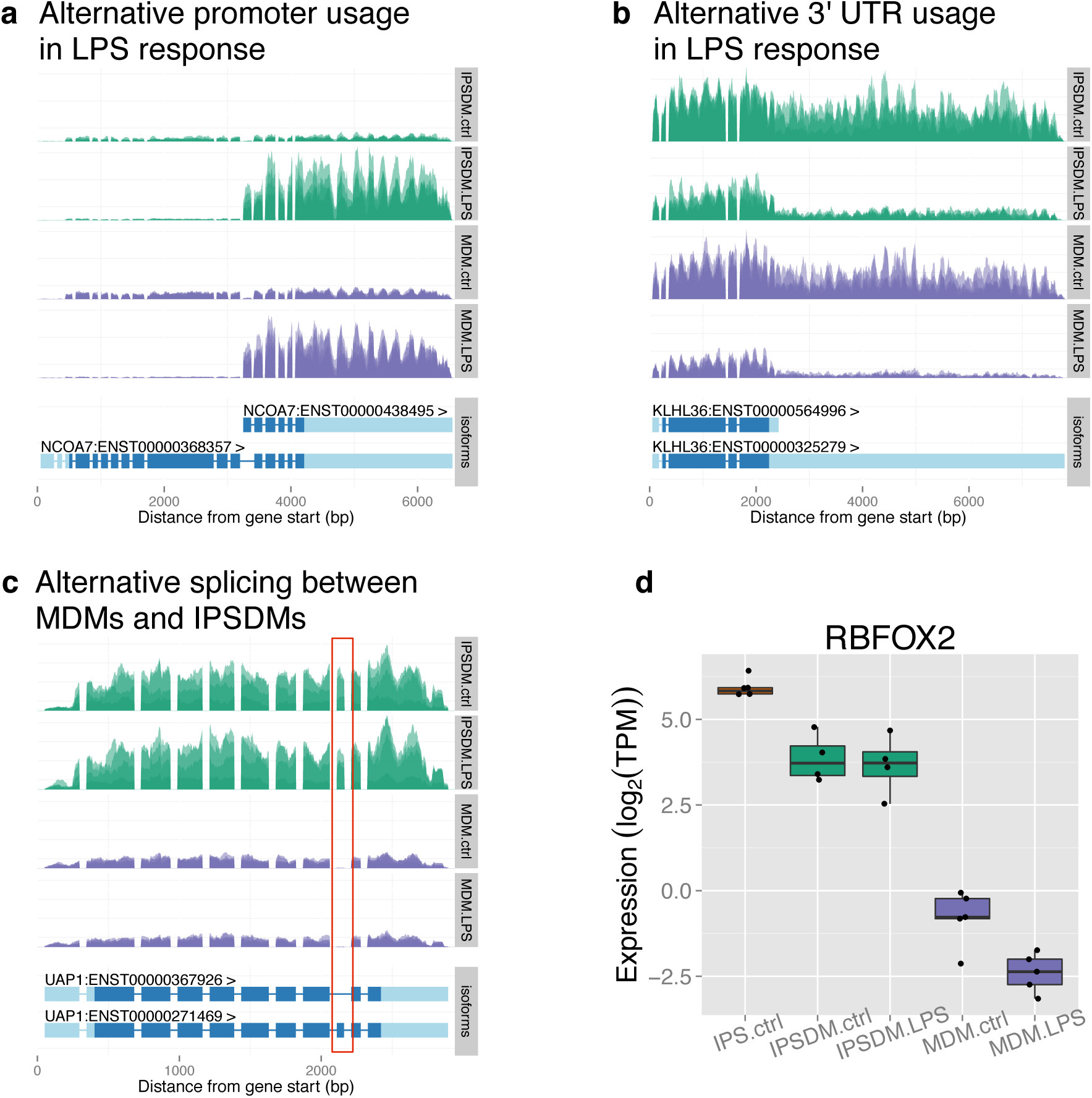
Examples of alternative transcript usage. Each plot shows normalised read depth across the gene body in IPSDMs (green) and MDMs (purple) with gene structure in the panel beneath each plot. Introns have been compressed relative to exons to facilitate visualisation. **(a)** Example of alternative promoter usage in LPS response. **(b)** Examples of 3′ UTR shortening in LPS response. **(c)** Example of alternative splicing between MDMs and IPSDMs. The alternatively spliced exon is marked with the red rectangle. **(d)** Expression of RBFOX2 gene in iPS cells, IPSDMs and MDMs.

We also observed widespread alternative 3′ end usage both in the LPS response as well as between MDMs and IPSDMs (Fig. 4c-d). In contrast to alternative promoters, most of the 3′ end events only changed the length of the 3′ UTR and not the coding sequence (Fig. 4d). Changes in 3′ UTR usage were strongly asymmetric: relative expression of transcripts with long 3′ UTRs was higher in IPSDMs compared to MDMs and it was also higher in unstimulated cells compared to stimulated cells (Fig. 4e, Fig. 5b). Notably, the observed pattern of decreased relative expression of transcripts with long 3′ UTRs from unstimulated IPSDMs to stimulated MDMs corresponded to the second principal component of relative transcript expression (Fig. 4a). Consistent with this observation, we found that genes with 3′ UTR events had higher than expected weights in PC2 (p < 2.2*10-16, chi-square goodness-of-fit test), (Supplementary Fig. S5) indicating that they are partially responsible for the observed distribution of samples along PC2. We found no convincing pathway or Gene Ontology enrichment signal in genes with 3′ UTR events.

Finally, we detected only a small number of alternative splicing events influencing middle exons, most of which occurred between MDMs and IPSDMs rather than in the LPS response (Fig. 4c-d). Three of the events with largest changes in proportion affected cassette exons in genes UAP1 (Fig. 5c), CTTN and CLSTN1 (Supplementary Fig. S5). Interestingly, the inclusion of these exons has previously been shown to be regulated by RNA-binding protein RBFOX2 that was significantly more highly expressed in IPSDMs (Fig. 5c)^19,20^.

## Discussion

In this study, we used high-depth RNA-Seq to investigate transcriptional similarities and differences between human monocyte and iPS-derived macrophages. Our principal findings are that relative to differences between MDMs and iPS cells, the transcriptomes of naïve and LPS stimulated MDMs and IPSDMs are broadly similar both at the whole gene and individual transcript levels. Although we have only examined steady-state mRNA levels, conservation of transcriptional response to LPS implies that the major components of signalling on protein level that coordinate this response must be similarly conserved. We did, however, also observe intriguing differences in expression in specific sets of genes, including those involved in tissue remodelling, antigen presentation and neutrophil recruitment, suggesting that IPSDMs might possess some phenotypic differences from MDMs. Our analysis also revealed a rich diversity of alternative transcription changes suggesting widespread fine-tuning of regulation in macrophage LPS response.

There are a number of possible explanations for the observed differences in transcription between MDMs and IPSDMs. Although our IPSDM samples could include minority populations of other cell types these were not obvious and all of our IPSDM samples were highly pure (92-99% CD14+) (Supplementary Fig. S7, Supplementary Table S6). Similarly, our MDM differentiation protocol routinely yields 90-95% pure cells, excluding contamination as the major source of these differences. Secondly, it is possible that some of the differences between IPSDMs and MDMs could be due to genetic differences between the donors. However, since all our IPSDM and MDM donors were sampled randomly from the same population, strong clustering of IPSDM and MDM samples on the heatmap (Supplementary Figure S1) suggests that genetics is not a major source of differences between these cell types. To address this quantitatively, we reanalysed an independent RNA-Seq data from 84 British individuals^21^. We found only a median of three differentially expressed genes between any two random samples of 4 and 5 individuals (Supplementary Fig. S8). This suggests that only a small fraction of the differences between MDMs and IPSDMs are likely to be due to genetic differences.

Alternatively, IPSDMs could show incomplete differentiation from iPS cells. Consistent with this hypothesis, genes that were more highly expressed in IPSDMs were often also expressed in iPS cells (Fig. 2c, Supplementary Fig. S9a) and large fraction of these genes had very low absolute expression in IPSDMs (Supplementary Fig. S9b). Furthermore, the promoters of the upregulated genes were highly enriched for repressive H3K27me3 histone marks in CD14+ monocytes^22^ (Supplementary Fig. S9c), suggesting that these genes become silenced prior to monocyte-macrophage differentiation *in vivo* and may not have been completely silenced in IPSDMs. Finally, IPSDMs might share some features with tissue resident macrophages that are developmentally and phenotypically distinct from MDMs^23–26^. In support of that, higher expression of tissue remodelling and neutrophil recruitment genes has previously been associated with tissue and tumour associated macrophages^27–30^. On the other hand, higher expression of antigen presentation genes in MDMs is consistent with the specialised role of monocyte-derived cells in immune regulation and antigen presentation^30–32^. This is consistent with a previous study suggesting a shared developmental pathway between IPSDMs and foetal macrophages^33^. Nevertheless, it is likely that the exact characteristics of IPSDMs can be shaped by the addition of cytokines and other factors during differentiation and this could be an important area for further exploration.

In addition to showing that LPS response was broadly conserved between MDMs and IPSDMs both on gene and transcript level, we also identified hundreds of individual alternative transcription events, highlighting an important, but potentially overlooked, regulatory mechanism in innate immune response. A small number of the events have known functional consequences. For example, the LPS-induced short isoform of the NCOA7 (Fig. 5a) gene is known to be regulated by Interferon β-1b and it is suggested to protect against inflammation-mediated oxidative stress^34^ whereas the long isoform is a constitutively expressed coactivator of estrogen receptor^35^. Similarly, the two isoforms of the OSBPL1A gene (Supplementary Fig. S4) have distinct intracellular localisation and function^36^ while the LPS-induced short transcript of the OSBPL9 gene (Supplementary Fig. S4) codes for an inhibitory isoform of the protein^37^. Thus, alternative promoter usage has the potential to significantly alter gene function in LPS response and these changes can be missed in gene level analysis.

Widespread shortening of 3′ UTRs has previously been observed in proliferating cells and cancer as well as activated T-cells and monocytes^38,39^. The functional consequences of 3′ UTR shortening are unclear, but extended 3′ UTRs are often enriched for binding sites for miRNAs or RNA-binding proteins that can regulate mRNA stability and translation efficiency^38,40^. The role of miRNAs in fine-tuning immune response is well established^41^. Furthermore, interactions between alternative 3′ UTRs and miRNAs have recently been implicated in the brain^42,43^. Therefore, it might be interesting to explore how 3′ UTR shortening affects miRNA-dependent regulation in LPS response.

In summary, we have performed an in depth comparison of an iPS derived immune cell with its primary counterpart. Our study suggests that iPS-derived macrophages are potentially valuable alternative models for the study of innate immune stimuli in a genetically manipulable, non-cancerous cell culture system that is likely superior to commonly used THP-1 monocytic cell line. The ability to readily derive and store iPS cells potentially enables in-depth future studies of the innate immune response in both healthy and diseased individuals. A key advantage of this model will be the ability to study the impact of human genetic variation, both natural and engineered, in innate immunity.

## Methods

### Samples

Human blood for monocyte-derived macrophages was obtained from NHS Blood and Transplant, UK and all experiments were performed according to guidelines of the University of Oxford ethics review committee. All IPSDMs were differentiated from four iPS cell lines: CRL1, S7RE, FSPS10C and FSPS11B. CRL1 iPS cell line was originally derived from a commercially available human fibroblast cell line and has been described before^44^. S7RE iPS cell line was derived as part of an earlier study from our lab^45^. FSPS10C and FSPS11B iPS cell lines were derived as part of the Human Induced Pluripotent Stem Cell Initiative. All iPS cell work were carried out in accordance to UK research ethics committee approvals (REC No. 09/H306/73 & REC No. 09/H0304/77).

### Cell culture and reagents

IPS cells were grown on Mitomycin C-inactivated mouse embryo fibroblast (MEF) feeder cells in Advanced DMEM F12 (Gibco) supplemented with 20% KnockOut Serum Replacement (Gibco, cat no 10828-028), 2mM L-glutamine, 50 IU/ml penicillin, 50 IU/ml streptomycin and 50 μM 2-mercaptoethanol (Sigma M6250) on 10 cm tissue-culture treated dishes (Corning). The medium was supplemented with 4 ng/ml rhFGF basic (R&D) and changed daily (10 ml per dish). Prior to passage, the cells were detached from the dish with 1:1 solution of 1 mg/ml collagenase and 1mg/ml dispase (both Gibco). Human M-CSF producing cell line CRL-10154 was obtained from ATCC. The cells were grown in T150 tissue culture flasks containing 40 ml of medium (90% alpha minimum essential medium (Sigma), 10% FBS, 2mM L-glutamine, 50 IU/ml penicillin, 50 IU/ml streptomycin). On day 9 the supernatant was sterile-filtered and stored at −80°C.

IPS cells were differentiated into macrophages following a previously published protocol consisting of three steps: i) embryoid body (EB) formation, ii) production of myeloid progenitors from the EBs and iii) terminal differentiation of myeloid progenitors into mature macrophages^9^.

For EB formation, intact iPS colonies were separated from MEFs using collagenase-dispase solution, transferred to 10 cm low-adherence bacteriological dishes (Sterilin) and cultured in 25 ml iPS media without rhFGF for 3 days. Mature EBs were resuspended in myeloid progenitor differentiation medium (90% X-VIVO 15 (Lonza), 10% FBS, 2mM L-glutamine, 50 IU/ml penicillin, 50 IU/ml streptomycin and 50 μM 2-mercaptoethanol (Sigma M6250), 50 ng/ml hM-CSF (R&D), 25 ng/ml hIL-3 (R&D)) and plated on 10 cm gelatinised tissue-culture treated dishes. Medium was changed every 4-7 days. After 3-4 weeks, floating progenitor cells were isolated from the adherent EBs, filtered using a 40 μm cell strainer (Falcon) and resuspended in macrophage differentiation medium (90 % RPMI 1640, 10% FBS, 50 IU/ml penicillin and 50 IU/ml streptomycin) supplemented with 20% supernatant from CRL-10154 cell line. Approximately 7×10^5^ cells in 15 ml of media were plated on a 10 cm tissue-culture treated dish and cultured for 7 days until final differentiation. We observed that using supernatant instead of 100 ng/ml hM-CSF as specified in the original protocol^9^ did not alter macrophage gene expression profile (Supplementary Fig. S10).

Human monocytes (90-95% purity) were obtained from healthy donor leukocyte cones (corresponding to 450 ml of total blood) by 2-step gradient centrifugation^46,47^. The monocyte fraction in this type of preparation is on average 98% CD14^+^, 13% CD16^+^ by single staining. The isolated monocytes were cultured for 7 days in the same macrophage differentiation medium as IPSDMs. The same seeding density and tissue-culture treated plastic was used as for IPSDMs. Non-adherent contaminating cells were removed by vigorous washing before cell lysis at day 7.

On day 7 of macrophage differentiation, medium was replaced with either 10 ml of fresh macrophage medium (without M-CSF) or medium supplemented with 2.5 ng/ml LPS (*E. coli*). After 6 hours, cells were lifted from the plate using lidocaine solution (6 mg/ml lidocaine, PBS, 0.0002% EDTA), counted with haemocytometer (C-Chip) and lysed in 600 μl RLT buffer (Qiagen). All cells from a dish were used for lysis and subsequent RNA extraction. The cell counts for IPSDMs samples are given in Supplementary Table S7.

### RNA extraction and sequencing

RNA was extracted with RNeasy Mini Kit (Qiagen) according to the manufacturer’s protocol. After extraction, the sample was incubated with Turbo DNase at 37°C for 30 minutes and subsequently re-purified using RNeasy clean-up protocol. Standard Illumina unstranded poly-A enriched libraries were prepared and then sequenced 5-plex on Illumina HiSeq 2500 generating 20-50 million 75bp paired-end reads per sample. RNA-Seq data from six iPS cell samples was taken from a previous study^45^. Sample information together with the total number of aligned fragments are detailed in Supplementary Table S7.

### Flow cytometry analysis

Flow cytometry was used to characterise the IPSDM cell populations used in the experiments. Approximately 1×10^6^ cells were resuspended in flow cytometry buffer (D-PBS, 2% BSA, 0.001% EDTA) supplemented with Human TruStain FcX (Biolegend) and incubated for 45 minutes on ice to block the Fc receptors. Next, cells were washed once and resuspended in buffer containing one of the antibodies or isotype control. After 1 hour, cells were washed three times with flow cytometry buffer and immediately measured on BD LSRFortessa cell analyser. The following antibodies (BD) were used (cat no): CD14-Pacific Blue (558121), CD32-FITC (552883), CD163-PE (556018), CD4-PE (561844), CD206-APC (550889) and PE isotype control (555749). The data were analysed using FlowJo. The raw data are available on figshare (doi: 10.6084/m9.figshare.1119735).

### Data analysis

Sequencing reads were aligned to GRCh37 reference genome with Ensembl 74 annotations using TopHat v2.0.8b^48^. Reads overlapping gene annotations were counted using featureCounts^49^ and DESeq2^11^ was used to identify differentially expressed genes. Genes with FDR < 0.01 and fold-change > 2 were identified as differentially expressed. We used gProfileR R package to perform Gene Ontology and pathway enrichment analysis^50^. For conditional enrichment analysis of the genes differentially regulated in LPS response we used all LPS-responsive genes as the background set. All analysis was performed on genes classified as expressed in at least one condition (TPM > 2) except where noted otherwise. The bedtools suite was used to calculate genome-wide read coverage^51^. All downstream analysis was carried out in R and ggplot2 was used for figures.

To quantify alternative transcript usage, reads were aligned to Ensembl 74 transcriptome using bowtie v1.0.0^52^. Next, we used mmseq and mmdiff to quantify transcript expression and identify transcripts whose proportions had significantly changed^15,18^. For each transcript we estimated the posterior probability of five models (i) no difference in isoform proportion (null model), (ii) difference between LPS treatment and controls (LPS effect), (iii) difference between IPSDMs and MDMs (macrophage type effect), (iv) independent treatment and cell type effects (both effects), (v) LPS response different between MDMs and IPSDMs (interaction effect). We specified the prior probabilities as (0.6, 0.1, 0.1, 0.1, 0.1) reflecting the prior belief that most transcripts were not likely to be differentially expressed. Transcripts with posterior probability of the null model < 0.05 were considered significantly changed.

We used two-step analysis to identify alternative transcription events from alternative transcripts. First, to identify all potential alternative promoter, alternative splicing and alternative 3′ end events in each gene, we compared the most significantly changed transcript to the most highly expressed transcript of the gene (Supplementary Fig. S11). Next, we reanalysed the RNA-Seq data using exactly the same strategy as described above (bowtie + mmseq + mmdiff) but substituted Ensembl 74 annotations with the identified alternative transcription events. This allowed us to separate the events truly supported by the data from the ones that were identified only because they were on the same transcript with a causal event (Supplementary Fig. S11). Finally, we required events to change at least 10% in proportion between two conditions to be considered for downstream analysis. The code to identify alternative transcription events from two transcripts is implemented in the reviseAnnotations R package (https://github.com/kauralasoo/reviseAnnotations). Our event-based approach is similar to the one used by MISO^53^.

## Acknowledgements

This work was supported by the Wellcome Trust grant #098051. K.A. was supported by a PhD fellowship from the Mathematical Genomics and Medicine programme from the Wellcome Trust. We thank Kosuke Yusa and Mariya Chhatriwala for fruitful discussions on troubleshooting iPS cell culture.

## Author contributions

Contribution: K.A., S.M. and D.G. designed the experiment; K.A. and C.H. performed the IPSDM experiments; F.M.E. performed the MDM experiments; K.A. analysed the data; K.A., S.M., D.G., G.D., F.P. and S.G interpreted the results; K.A., D.G. and S.M. wrote the manuscript; D.G., G.D. S.G. and F.P. supervised the research. All authors discussed the results and commented on the manuscript.

## Accession codes

RNA-Seq data are available from the European Genome-phenome Archive (EGA) under accession EGAS00001000563.

